# sRNAs enriched in outer membrane vesicles of pathogenic *Flavobacterium psychrophilum* interact with immune genes of rainbow trout

**DOI:** 10.1101/2021.12.22.473952

**Authors:** Pratima Chapagain, Ali Ali, Destaaalem T. Kidane, Mary Farone, Mohamed Salem

## Abstract

Outer membrane vesicles (OMVs) released by gram-negative bacteria during host-pathogen interactions harbor cargos, such as DNA, RNA, toxins, and virulence factors. We hypothesized that sRNAs carried within OMVs of *Flavobacterium psychrophilum* interact with host immune genes and affect their expression. OMVs were isolated from *F. psychrophilum* and visualized using transmission electron microscopy (TEM). RNA-Seq datasets generated from whole-cell *F. psychrophilum* and their OMVs indicated enrichment of specific sRNAs in the OMVs compared to the parent cell. Fluorescent *in situ* hybridization (FISH) and confocal microscopy confirmed the expression of a randomly chosen sRNA.

Integrated RNA-Seq analyses of host transcriptome and bacterial sRNAs on day 5 post-infection of *F. psychrophilum*-resistant and -susceptible rainbow trout genetic lines revealed 516 protein-coding, 595 lncRNA, and 116 bacterial sRNA differentially expressed (DE) transcripts. Integrated and network analyses of these DE transcripts revealed immune genes targeted by bacterial sRNAs. On the top of these genes, an isoform encoding anaphase-promoting complex subunit 13 (ANAPC13_1) was highly upregulated and exhibited interaction and reciprocal expression with 21 DE sRNAs enriched in OMVs and/or located in pathogenicity islands (PAIs). *In vitro* treatment of the rainbow trout epithelial cell line RTgill-W1 with OMVs showed signs of cell autolysis accompanied by dynamic changes in expression of host genes when profiled 24h following treatment. The OMV-enriched sRNAs, *soFE013584* and *soFE002123*, showed high interactions with the protection of telomeres 1 gene (POT1); essential for chromosome stability and cellular viability. Modulation of the host gene expression following OMV-treatment, which favors elements from the phagocytic, endocytic, and antigen presentation pathways in addition to HSP70, HSP90, and cochaperone proteins, provided evidence for a potential role of OMVs in boosting the host immune response. In conclusion, our work identified novel microbial targets and inherent characteristics of OMVs that could open up new avenues of treatment and prevention of fish infections.

## Introduction

*Flavobacterium psychrophilum* causes Bacterial cold-water disease (BCWD), a severe infection that frequently affects salmonid species, including salmon and rainbow trout. BCWD is responsible for causing a considerable economic loss in freshwater aquaculture of salmonids [1]. Losses from this infection have been reported in many countries, including the United States, United Kingdom, Australia, and Japan [2]. Mortality rates from this disease vary, but rates as high as 90% have been reported [3]. *F. psychrophilum* is a psychrophilic, gram-negative bacterium typically found in cold freshwater [4]. After entering the host, likely through skin abrasions, *F. psychrophilum* spreads through the dermis, connective, and muscle tissues [1]. Rainbow trout (especially fry) infected by *F. psychrophilum* exhibit irregular swimming behavior, exophthalmia, darkened coloration, skeletal deformities, and visible deep necrotic lesions around the caudal peduncle [3]. The survivability of this pathogen under various environmental conditions and the unavailability of vaccines contribute to the severity of the disease [5].

Efforts have been made to create BCWD-resistant strains of rainbow trout. Recent approaches have correlated fish genetic variations to immune responsiveness and survival rates after challenges with *F. psychrophilum* [6–8]. Through a selective breeding program, the National Center for Cool and Cold-Water Aquaculture (NCCCWA) developed three genetic lines of rainbow trout: BCWD-susceptible, -control, and -resistant lines [6, 9]. The susceptible line has a very low survival rate (29.4%), the control line has an intermediate survival rate (54.6%), and the resistant line has undergone multiple generations of selection against *F. psychrophilum* challenge to enhance their survival rate up to 94% [6].

The molecular pathogenesis of *F. psychrophilum* is still not yet completely understood. Many factors, including biofilm formation, secretion systems, and virulence factors such as exotoxins, proteases, and adhesins, are thought to play a role in the pathogenesis of this bacterium [1, 10]. *F. psychrophilum*, similar to other gram-negative bacteria, produce small (< 300 nm in diameter) spherical particles derived from the bacterial outer membrane, commonly known as outer membrane vesicles (OMVs) [11]. Many pathogenic bacteria rely on OMVs to deliver virulence factors to their host [12, 13], and OMV-mediated delivery has been considered an essential mechanism in host-pathogen interactions. For example, OMVs of *Porphyromonas gingivalis* contain a group of proteases, constituting the virulence factor “gingipains”, which degrade cytokines and ultimately downregulate the immune response of host cells [14]. Similarly, different cargos, such as DNA, RNA, and cytosolic proteins within OMVs, can be transported to the host cell when OMVs fuse with host cell membranes or bind to the host cell receptors [15–17]. sRNAs within OMVs of *Pseudomonas aeruginosa* have been shown to reduce the host immune response [16].

Bacterial sRNAs are non-protein-coding molecules with a length typically ranging from 50 to 500 nucleotides and have essential roles in post-transcriptional gene regulation [18, 19]. sRNAs usually originate from the untranslated regions of the bacterial genome but can also be acquired by horizontal gene transfer [18, 20]. Many sRNAs act as complementary antisense sequences to mRNAs [21], and their function resembles that of the eukaryotic microRNAs (miRNAs). They cause translational repression by directly hybridizing with mRNAs to promote transcript degradation [22]. These molecular mechanisms allow sRNAs to play a role in bacterial pathogenicity. For example, a few studies have suggested that the network of bacterial genes in biofilm formation is controlled by sRNAs [23, 24]. A recent study in *P. aeruginosa* has shown the involvement of sRNAs in repressing the translation of genes involved in quorum sensing by binding to the ribosome binding sites of mRNA targets [25]. A study on *Staphylococcus aureus* reported the participation of sRNAs in regulating cellular responses to signal molecules [26]. Another study reported that entry of bacterial sRNAs inside host cells triggers a host gene-bacterial sRNAs interaction resulting in degradation of host immune genes [27].

In this study, OMVs were isolated from *F. psychrophilum* (strain CSF-259-93) broth culture on day 8 of bacterial growth. The parent bacteria and OMVs were subjected to RNA sequencing, and then the transcriptomic data were used for sRNAs prediction. We identified sRNAs specific and/or enriched in OMVs by comparing sRNA expression levels in the parent cells versus OMVs. Further, *in vivo* investigated the interactions among bacterial sRNAs within *F. psychrophilum* OMVs and rainbow trout immune genes. Dual RNA-Seq of host and pathogen revealed that bacterial sRNAs were reciprocally expressed with their target immune genes on day 5 following infection of selectively bred resistant-, and susceptible-line rainbow trout. These results suggest a role for bacterial sRNAs during host-pathogen interactions, facilitating bacterial invasion/growth. We then *in vitro* investigated the effect of OMVs on RTgill-W1 host cells and the interaction between bacterial sRNA and host genes. OMVs caused host cell lysis; however, immune response genes expression was boosted at the early phase of OMV treatment (24h post-treatment).

## Results

### OMVs extraction and TEM

OMVs were isolated from bacterial cells for downstream RNA sequencing and prediction of bacterial sRNAs. The experimental design is shown in Fig. 1. *F. psychrophilum* colonies grown on Tryptone Yeast Extracts (TYEs) agar plates were observed as bright yellow colonies after 5 days of incubation at 15°C. In broth culture, bacterial log phase growth was observed from day 5 until day 11 (Fig. 2a). Isolated *F. psychrophilum* OMVs were observed by transmission electron microscopy (TEM) as spherical-shaped particles with an average diameter of 50-100 nm (Fig. 2b), whereas *F. psychrophilum* was observed as having rod-shaped morphology with a size of approximately 3-5 μm (Fig. 2c).

**Fig. 1:**
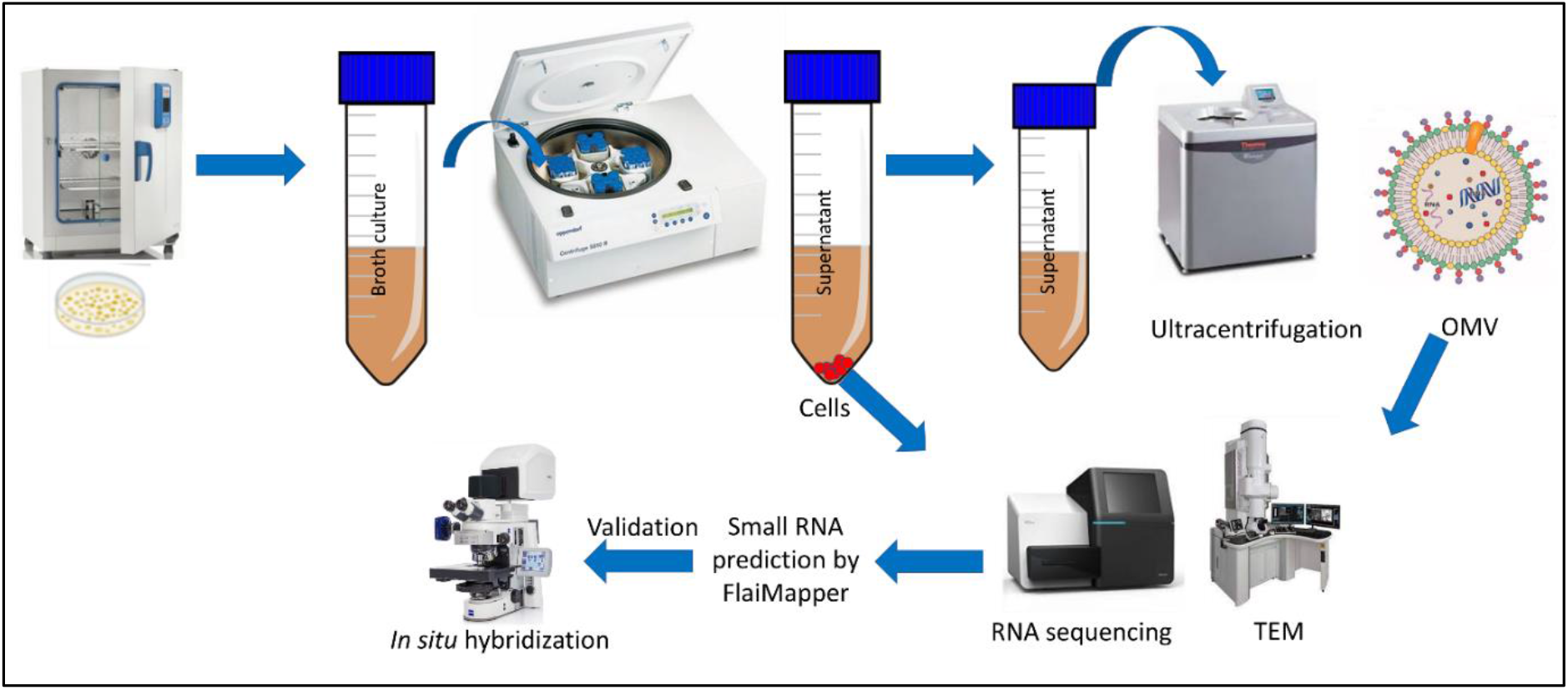
Overview of the experimental design to predict and validate bacterial small RNAs. Frozen stock cultures of *F. psychrophilum* were cultured on TYEs agar, and the plate was incubated at 15°C for one week. *F. psychrophilum* colonies isolated from TYEs agar plate were transferred to TYEs broth. For OMV isolation, broth culture tubes were centrifuged to pellet the bacterial cells, and the supernatant was collected and filtered to remove any remaining bacterial cells. The filtrate was then subjected to ultracentrifugation to pellet the OMVs. *F. psychrophilum* cells and OMVs were visualized using TEM, followed by RNA sequencing on an Illumina MiSeq platform. Small RNAs were predicted using FlaiMapper, and validated by FISH.

**Fig. 2:**
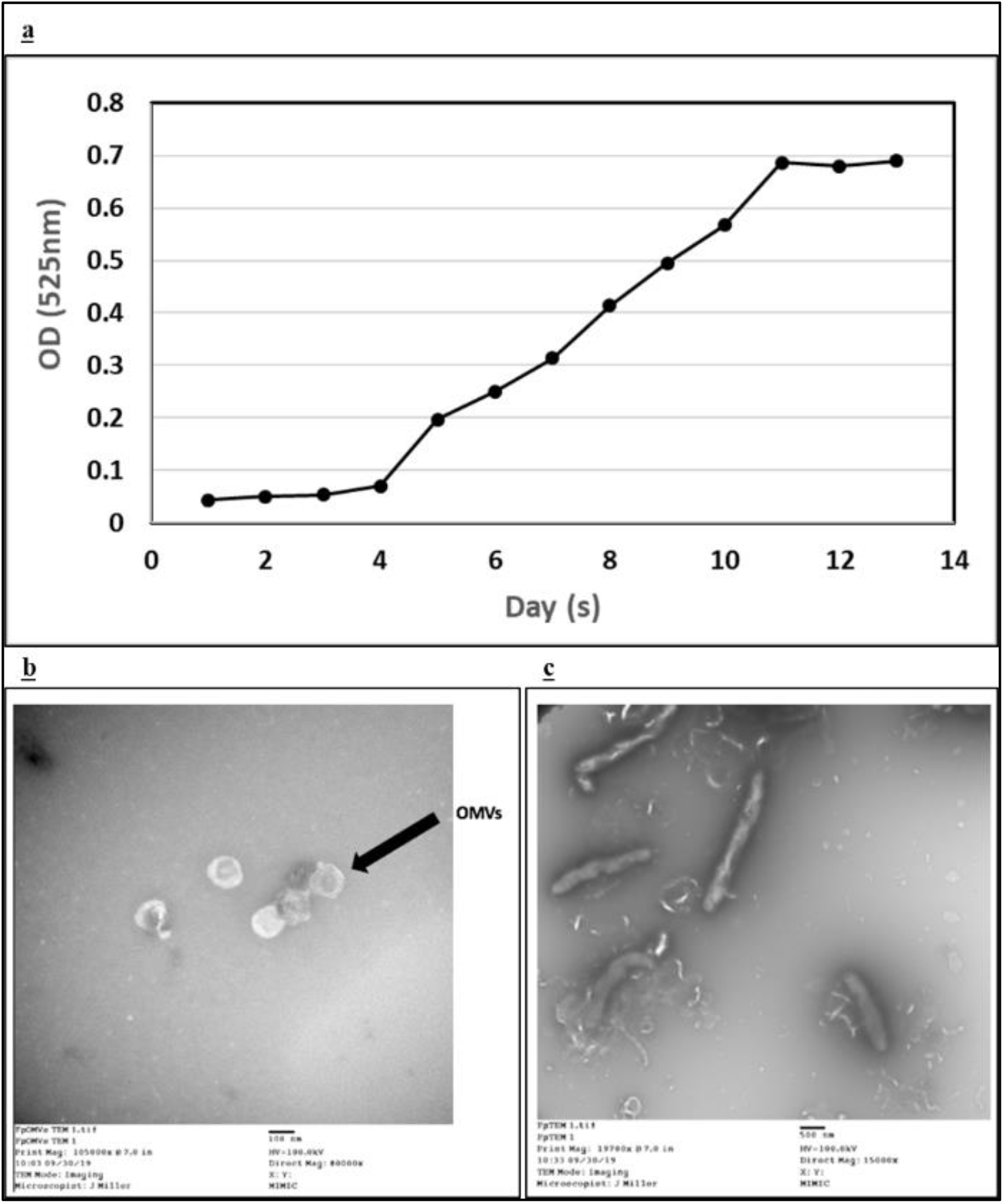
**a)** Measurement of *F. psychrophilum* density by measuring OD in broth culture, the log-phase was observed between days 5 and 11. **b)** Transmission Electron Microscopy (TEM) of *F. psychrophilum* OMVs. OMVs appeared as spherical-shaped particles, and the black arrow points to the bilayered OMVs’ spherical structure. **c)** TEM of *F. psychrophilum*, which appeared as a rod-shaped structure.

### RNA-Seq identifies sRNAs enriched in OMVs

Total RNA isolated from whole-cells and OMVs was sequenced, yielding 60,352,578 and 55,722,742 sequence reads, respectively. High-quality reads from whole-cell and OMVs were separately mapped to the *F. psychrophilum* reference genome, where 80.0% and 80.3% read mapping rates were achieved, respectively. FlaiMapper [28] was used to predict small noncoding RNAs from the alignment files. In total, 118,876 and 134,518 noncoding transcripts were predicted from the whole-cells and OMVs, respectively. Transcripts longer than 500 bp were filtered out, yielding 118,305 and 133,815 predicted sRNAs from the whole-cells and OMVs, respectively. The sRNA length ranged from 87 to 499 nucleotides (Average 100.4 nt). The majority of the predicted sRNAs (98.9%) were ≤100 bp in length. The list of predicted sRNAs with their genomic location is included in Additional file 1 (Tables S1 & S2). The sRNA *soFE128978* was randomly selected for validation and visualization by confocal microscopy. The Cy3 fluorophore was used to visualize the sRNA. Fluorescent *in situ* hybridization and confocal imaging confirmed the expression of the sRNA within *F. psychrophilum* (Fig. 3). The red color in *F. psychrophilum* using an antisense probe and no fluorescence in *F. psychrophilum* using a non-sense probe (negative control) validates the results.

**Fig. 3:**
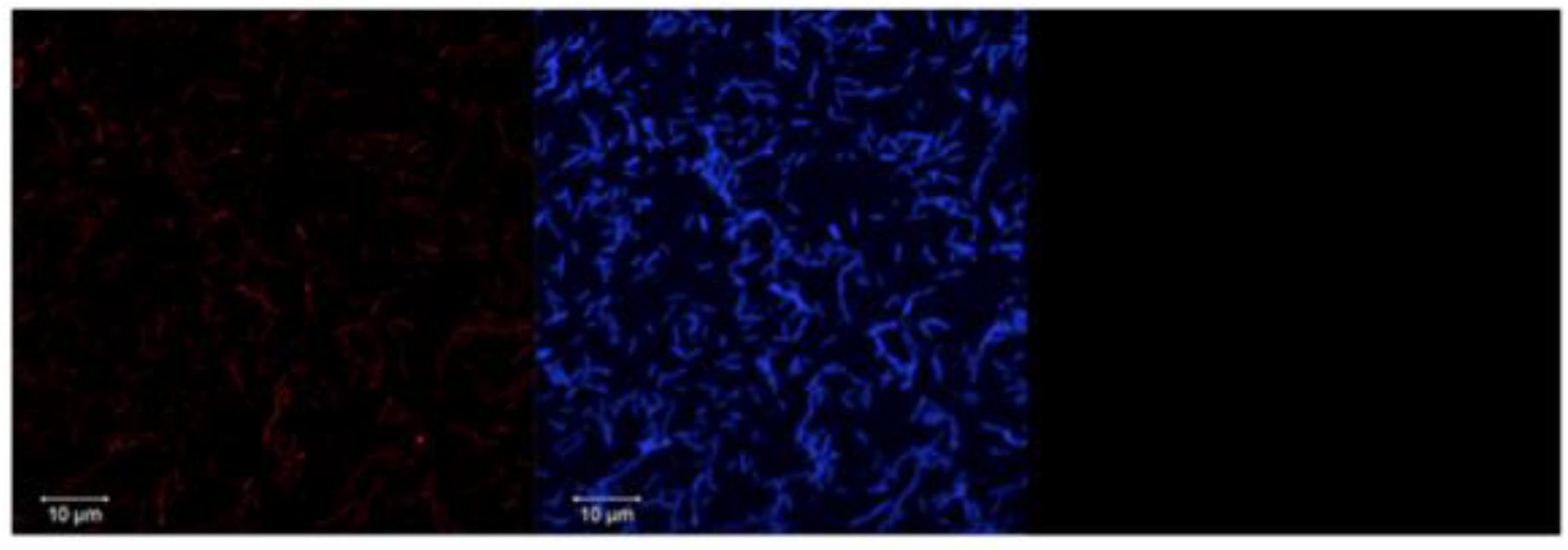
Confocal micrograph of FISH stained *F. psychrophilum* cells using Cy3-labeled *soFE128978* probe (Red) on the left panel. No fluorescence was observed in *F. psychrophilum* using a non-sense probe on the right panel (negative control) and *F. psychrophilum* cell stained by DAPI in the center panel.

RNA-Seq of the whole-cell and OMVs revealed 11,015 and 10,437 sRNAs, respectively, with Transcripts Per Million (TPM) expression level greater than 0.5. Of them, 5,975 sRNAs in OMVs and 5,947 sRNAs in the whole cell had TPM >1. The expression value of sRNAs in OMVs and the whole cell are included in Additional file 1 (Tables S1 & S2). sRNAs were classified as OMV-specific when OMV sRNAs share less than 90% sequence overlap with sRNAs in the whole cell. Otherwise, they were considered “common” sRNAs. We reported the sRNAs classification in Additional file 1 (Tables S3 & S4). In total, 108 OMV-specific sRNAs with TPM > 0.5 were identified (Table 1 and Additional file 1; Table S3). Also, sRNAs were classified as OMV-enriched if their TPM expression was at least two times higher than the whole cell (log_2_ TPM ratio ≥ 1). 17.23% of the common sRNAs were OMV-enriched (Table 2 and Additional file 1; Table S4). The OMV-specific or -enriched sRNAs could be explained by the selective inclusion of sRNAs in the OMV, which suggests potential roles of these sRNAs in microbe-host interaction. Consistent with our data, a previous study reported enrichment of sRNAs in *P. aeruginosa* OMVs [16].

**Table 1:** A subset of OMV-specific sRNAs with TPM > 1.5; PAI denotes sRNAs located in Pathogenicity Islands.

| sRNA | Start | End | TPM | PAI |
| --- | --- | --- | --- | --- |
| soFE129980 | 607022 | 607121 | 829.95 | Yes |
| soFE081095 | 2583062 | 2583161 | 13.10 | No |
| soFE023505 | 490867 | 490966 | 7.14 | No |
| soFE113400 | 2080493 | 2080592 | 5.60 | No |
| soFE014089 | 490862 | 490961 | 5.44 | No |
| soFE024485 | 490852 | 490951 | 2.27 | No |
| soFE068150 | 939025 | 939124 | 2.16 | No |
| soFE038667 | 2656387 | 2656486 | 2.11 | No |
| soFE132118 | 1665123 | 1665222 | 1.90 | Yes |
| soFE005824 | 1180568 | 1180667 | 1.86 | No |
| soFE017951 | 924082 | 924181 | 1.78 | No |
| soFE070737 | 209041 | 209140 | 1.58 | No |
| soFE071960 | 1201207 | 1201306 | 1.52 | No |
| soFE075699 | 1480007 | 1480106 | 1.52 | No |

**Table 2:** A subset sRNAs exhibiting enriched expression in the OMVs compared to the whole-cell. Log_2_ ratio is the logarithmic value obtained by dividing the TPM expression of each sRNA in OMVs by that of the whole cell. PAI denotes sRNAs located in Pathogenicity Islands.

| sRNA | Start | End | TPM (OMV) | TPM (cell) | log<sub>2</sub> ratio | PAI |
| --- | --- | --- | --- | --- | --- | --- |
| soFE000222 | 2080716 | 2080815 | 608.38 | 0.96 | 9.32 | No |
| soFE000671 | 2080722 | 2080821 | 199.97 | 0.96 | 7.71 | No |
| soFE000141 | 2080709 | 2080808 | 743.95 | 3.80 | 7.61 | No |
| soFE001158 | 2080734 | 2080833 | 18.32 | 0.11 | 7.41 | No |
| soFE000208 | 2080699 | 2080798 | 822.14 | 7.66 | 6.75 | No |
| soFE000754 | 2080729 | 2080828 | 43.14 | 0.66 | 6.03 | No |
| soFE017190 | 490892 | 490991 | 12.79 | 0.69 | 4.21 | No |
| soFE017916 | 490900 | 490999 | 12.94 | 0.77 | 4.07 | No |
| soFE031643 | 490880 | 490979 | 8.79 | 0.59 | 3.91 | No |
| soFE000238 | 2580160 | 2580259 | 958.52 | 66.57 | 3.85 | No |
| soFE000237 | 2092686 | 2092785 | 958.51 | 66.58 | 3.85 | No |
| soFE000236 | 1662630 | 1662729 | 958.54 | 66.58 | 3.85 | Yes |
| soFE005377 | 2046610 | 2046709 | 6.45 | 1.74 | 1.89 | No |
| soFE036061 | 2305986 | 2306085 | 0.54 | 0.15 | 1.88 | No |
| soFE007845 | 2046631 | 2046730 | 7.30 | 1.99 | 1.87 | No |
| soFE010415 | 862382 | 862481 | 1.24 | 0.40 | 1.65 | No |

### sRNAs within pathogenicity islands (PAIs)

Ten genomic islands were predicted in *F. psychrophilum* by at least one computational method (Fig. 4). The full output table from the IslandViewer showing genomic island size, location, and annotation is included in Additional file 2 (Table S5). 863 sRNAs with expression level TPM > 1 were predicted in the PAIs (Additional file 2; Table S6). For instance, the OMV-specific sRNA *soFE129980* was located within a PAI spanning the genomic region 583,837-607,648. Other genes within this PAI included transposase, integrase, universal stress protein UspA, and transcription termination/anti-termination protein NusA (Fig. 4). Also, we identified an OMV-specific sRNA, *soFE132118*, spanning the genomic region 1,660,689-1,665,551. Due to their pathogenic potential, we sought to investigate the interaction of OMV-specific sRNAs (Table 1) with immune-related target genes. Marancik et al. [6] identified 2,633 putative immune relevant genes by GO and manual annotations of the first draft of the trout genome [29]. We remapped these sequences to a newer reference of the rainbow trout genome. Sequences were mapped to 2,101 genomic loci. We selected the longest transcript generated from each locus to investigate the OMV sRNA-host immune gene interactions. Fourteen sRNAs (Table 1) exhibited computationally predicted interaction with 69.6% of the host immune transcripts (Additional file 2; Table S7). For instance, sRNA *soFE129980* targeted 1,117 of the host immune-relevant genes; of them, 151 genes exhibited very strong interactions (ndG cutoff −0.2). These results suggest a potential role for the OMV-sRNAs in mediating host-pathogen interactions to facilitate pathogen invasion/growth.

**Fig. 4:**
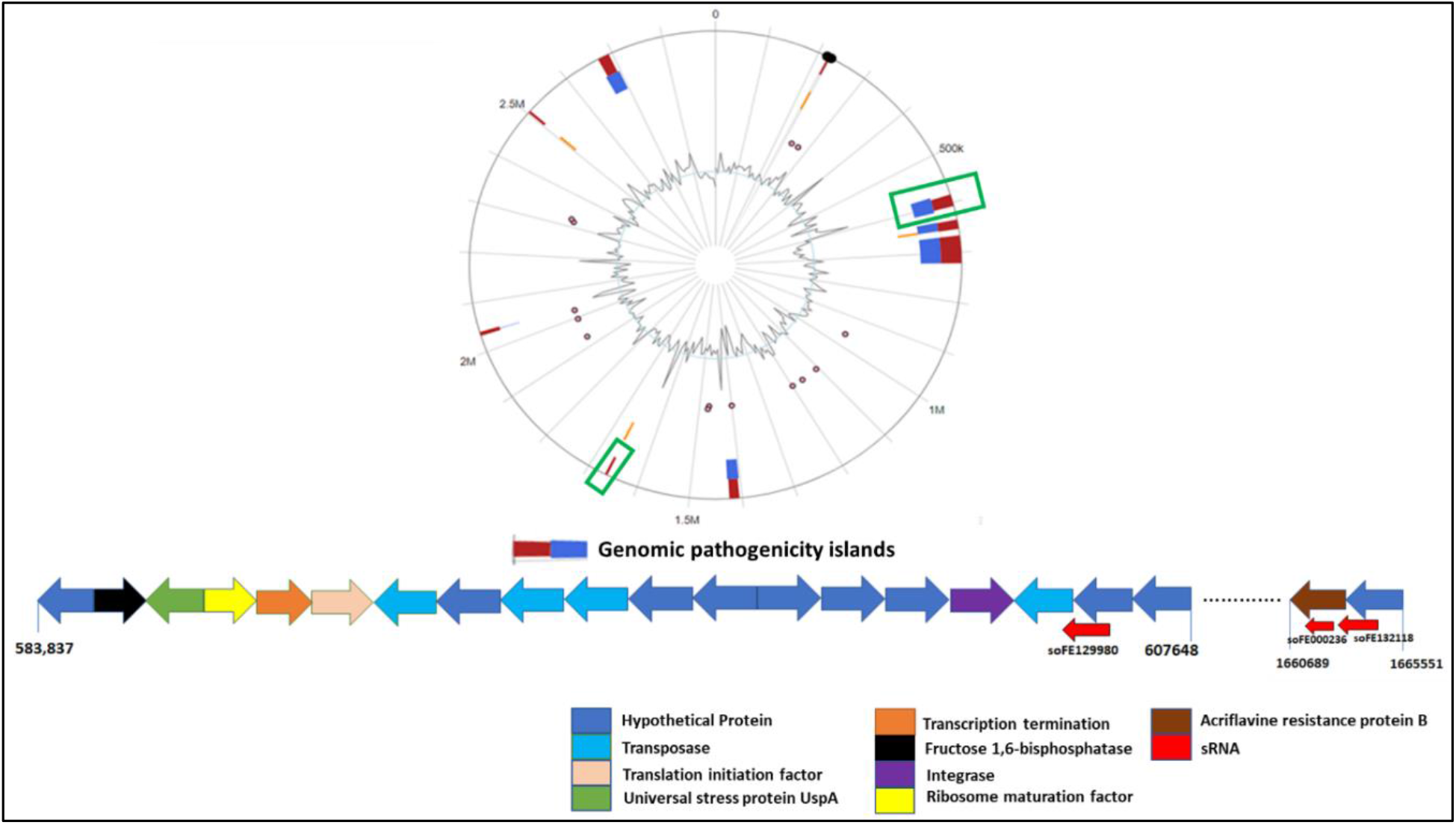
Genomic pathogenicity islands (clusters of genes that play a role in microbial adaptability) of *F. psychrophilum* predicted by either IslandPath-DIMOB alone (Blue) or integrated methods (at least two methods, Red). The green windows indicate the location of 3 sRNAs, *soFE129980, soFE132118* and *soFE000236*, within genomic pathogenicity islands. Color-coded arrows, representing the physical location of genes within PAIs, are annotated at the bottom of the figure, where the red arrows indicate sRNAs.

### Dual RNA-Seq of host and pathogen on day 5 post-infection of fish from resistant and susceptible genetic lines

In this study, we aimed to investigate host-pathogen interactions, *in vivo*, on day 5 following infection of rainbow trout with *F. psychrophilum*. In particular, we sought to investigate the potential role of bacterial sRNAs in disease susceptibility and identify host immune-related genes and lncRNAs that are likely targeted by these sRNAs. For this purpose, we used fish collected from selectively bred resistant (ARS-Fp-R) and susceptible (ARS-Fp-S) genetic lines challenged with *F. psychrophilum* as previously described in [6]. Total RNA was sequenced from 8 pooled samples for host mRNAs, lncRNAs, and bacterial sRNAs (see Methods section). RNA-Seq from infected resistant and susceptible genetic lines yielded a total of 372,100,715 raw sequence reads (average of 46,512,589 reads/sample) with ~12.1 depth of coverage. Sequence reads were trimmed/filtered to generate 371,959,119 high-quality reads (average of 46,494,890 reads/sample). To identify DE transcripts, high-quality reads were separately mapped to the reference genomes of the host and pathogen (Fig. 5a). A total of 285,744,993 (76.8%) trimmed reads were mapped to the rainbow trout genome. In our previous studies, 59.9% of the total RNA-sequence reads (518,881,838) were mapped to the trout references [6, 7]. Additionally, 4,729,437 reads (1.27%) were mapped to the *F. psychrophilum* reference genome. Notably, 99.4% of the *F. psychrophilum* mapped reads were generated from the susceptible genetic line. A higher bacterial load was previously reported in the susceptible line when compared to the resistant line [6]. Normalized gene expression was used to account for differences across samples by converting raw count data to Transcripts Per Kilobase Million (TPM) format. Quality and mapping statistics of sequence reads are given in Additional file 3; Table S8. A total of 516 (483 loci) protein-coding, 595 lncRNA, and 116 bacterial sRNA transcripts were DE with FDR < 0.05 and a minimum log_2_ fold change value ≥ 1 or ≤−1 (Fig. 5b & 5c and Additional file 3; Tables S9, S10 & S11). Of the DE protein-coding transcripts, 82 had immune-related functions, as we previously reported [6]. Most of the DE protein-coding (86.2%) and lncRNA transcripts (68.7%) were downregulated in the resistant line. In our previous study, 54 out of 83 lncRNAs (65.1%) were downregulated on day 5 post-infection [7]. Notably, 26.5% of the differentially regulated lncRNAs previously described in [7] were also identified in the current study, suggesting the potential role of these genes in resistance to BCWD. More information about DE transcripts is given in Additional file 3 (Tables S9, S10 & S11).

**Fig. 5:**
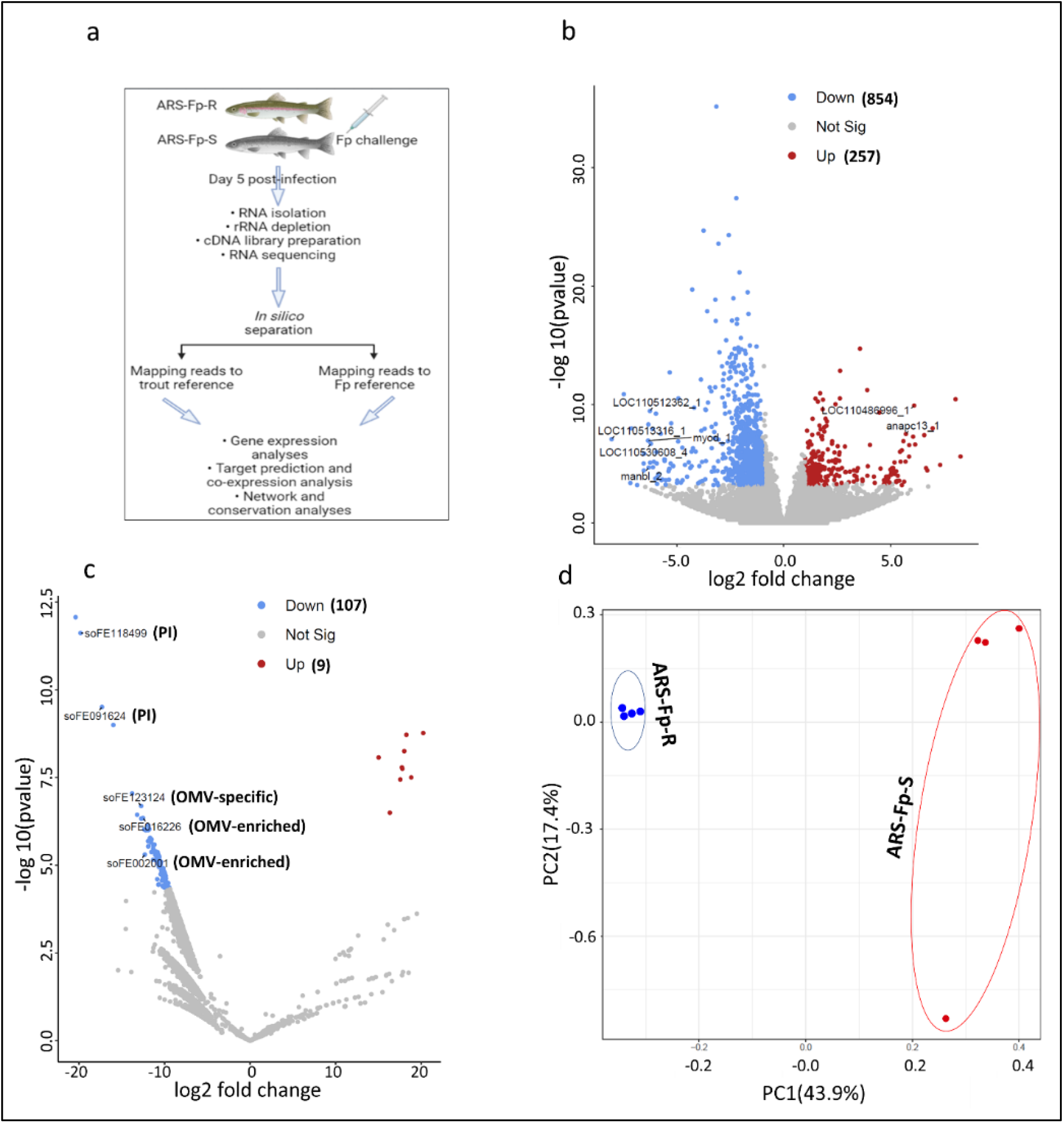
**a)** Whole-body dual RNA-seq. Fish were intraperitoneally injected with *F. psychrophilum* as previously described by [6]. Total RNA was isolated from fish collected on day 5 post-infection and processed for sequencing. Sequence reads were separated, *in silico*, by mapping to the rainbow trout and *F. psychrophilum* genomes to identify DE transcripts and hub genes during host-pathogen interactions. **b,c)** Volcano plots showing the host transcripts (mRNAs & lncRNAs) and bacterial sRNAs, respectively, differentially expressed on day 5 following *F. psychrophilum* infection in selectively bred, resistant-versus susceptible-line rainbow trout. The red dots represent the upregulated transcripts in the resistant line, whereas the blue dots represent the downregulated transcripts at FDR ≤ 0.05. Four sRNAs named on the figure, either located in the PAIs or enriched in OMVs, were among the most upregulated sRNAs in the susceptible genetic line. **d)** Principal component analysis of OMV-specific sRNAs obtained from 8 RNA-seq datasets generated from selectively bred, resistant- and susceptible-line rainbow trout on day 5 post-infection. Each round bullet represents a single RNA-seq dataset color-coded by a genetic line.

In order to gain insights into the implications and biological roles of bacterial sRNAs in fish following infection, we profiled their expression in resistant and susceptible genetic lines (Fig. 5c) and then investigated their expression correlation and interaction with the host genes (Fig. 6a and Additional file 3; Tables S12 & S13). Interestingly, a significant difference in the OMV-specific sRNA expression was detected between the genetic lines on day 5 following infection (Fig. 5d). A principal component analysis (Fig. 5d) showed that the primary axis accounts for ~44% of the variation. The pairwise comparison revealed 116 sRNAs with differential abundance between fish from the two genetic lines following infection (Additional file 3; Table S11). Of them, 28 DE sRNAs were OMV-specific. 107 sRNAs (92.2%) were upregulated in susceptible fish. A total of 71 DE sRNAs exhibited reciprocal expression correlation (R < −0.85) and interaction (ndG ≤ −0.2) with 143 DE host protein-coding transcripts (Additional file 3; Table S12). For instance, the OMV-specific sRNA *soFE085997*, followed by *soFE129182* and *soFE045225*, exhibited the highest potential of interaction with two isoforms encoding trichohyalin (Fig. 6b). The latter has an essential role in the proliferation and anti-apoptosis of human keratinocytes [30], which previously demonstrated a capacity to kill bacteria [31]. BCWD causes caudal lesions in rainbow trout, and thus the upregulation of trichohyalin in resistant fish may represent a defense mechanism against the pathogen. On the other hand, more DE bacterial sRNAs (98.3%) were correlated with DE host lncRNAs (70.6%) in expression and were more likely to interact with each other (ndG ≤ −0.2) (Additional file 3; Table S13). Four of the hosts’ lncRNAs showed expression correlation and interaction with the host trichohyalin isoforms and bacterial sRNAs (Fig. 6b and Additional file 3; Tables S13 & S14).

**Fig. 6:**
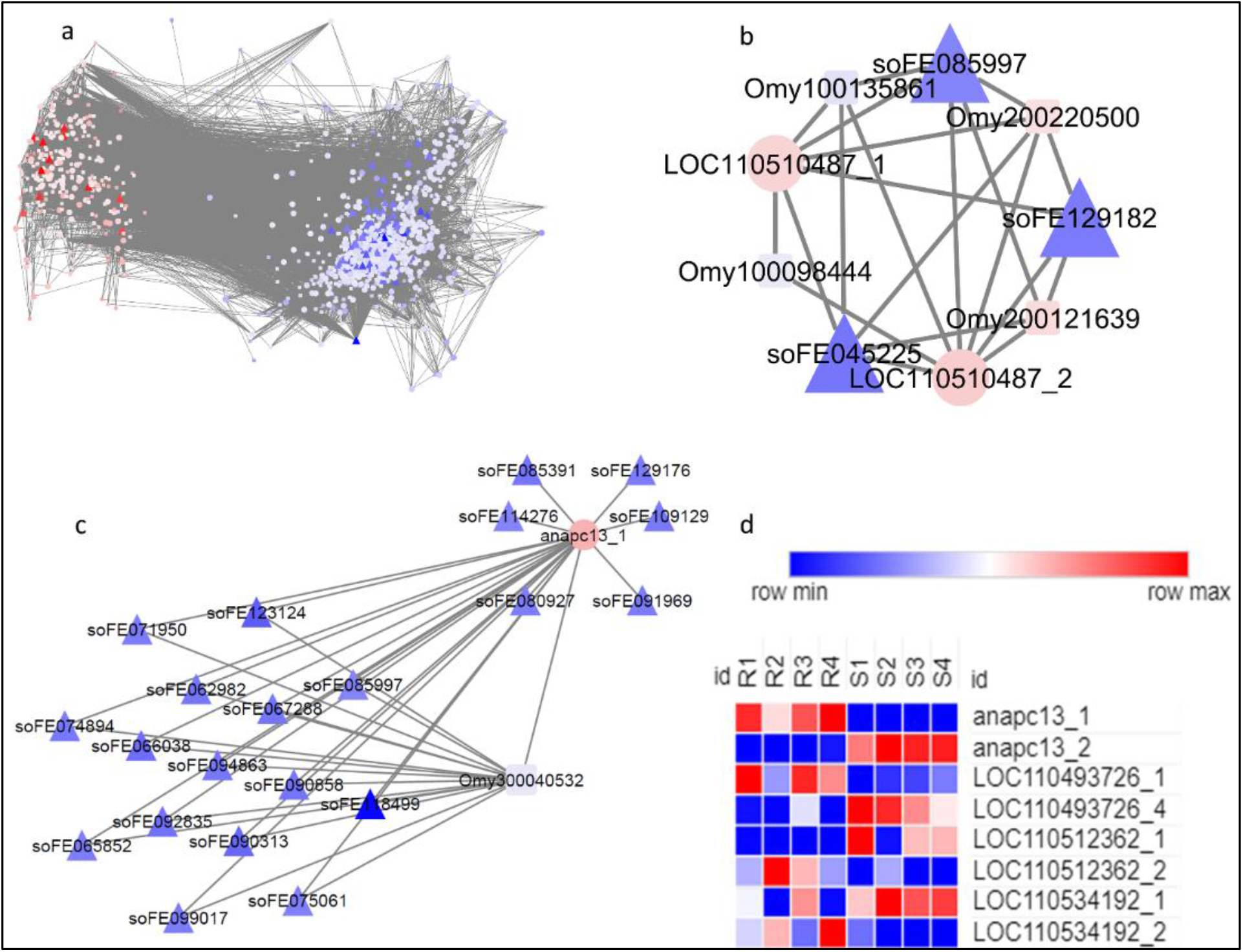
**a)** Gene expression network of DE bacterial sRNAs (triangular nodes) and DE host transcripts (mRNAs “circular nodes” and lncRNAs “rectangular nodes”) (R > 0.85 or <−0.85). DE transcripts are clustered into two groups based on their fold change, with the upregulated transcripts in the BCWD-resistant genetic line represented in red and the downregulated transcripts represented in blue. **b)** Three bacterial sRNAs (blue), including the OMV-specific sRNA *soFE085997*, exhibited reciprocal expression with two isoforms encoding trichohyalin (LOC110510487_1 & LOC110510487_2). **c)** ANAPC13_1 (red circle) was the top known downregulated transcript in susceptible fish on day 5 post-infection. ANAPC13_1 exhibited target interaction and negative correlation in expression with 21 sRNAs located in PAI or enriched in OMVs. LncRNA Omy300040532 likely mediates the interaction between most of these sRNAs and ANAPC13_1. **d)** Four isoforms, including ANAPC13_1, exhibited reciprocal expression with four other isoforms produced from the same genomic locus. “R1-4” refer to samples from the resistant genetic line, whereas “S1-4” refer to samples from the susceptible genetic line.

Among the differentially regulated protein-coding transcripts, 445 (86.2%) were downregulated in the resistant line (Additional file 3; Table S9). The list included genes coding for histone H3, MYOD protein, C-X-C motif chemokine 11, cathelicidin antimicrobial peptide, and hepcidin. Histone H3 (log_2_FC = −8.05) and MYOD (log_2_FC = −6.3) were at the top of the downregulated genes in the resistant line. Pathogens, such as bacteria, reprogram the host cells during infection through induction of histone H3 modifications, which modulate the host transcription machinery [32]. Notably, nine sRNAs enriched in the parent bacterial cells exhibited a strong reciprocal expression with histone H3. MYOD is a transcription factor involved in activating genes coding for muscle-specific proteins [33, 34]. Inactivity or reduced swimming activity has been recently reported to be associated with disease resistance in salmon [35]. In *Salmo salar*, myosin was upregulated in susceptible fish in response to sea lice infection [35]. MYOD did not directly show correlation/interaction with the bacterial sRNAs. However, MYOD showed a negative correlation (R = −0.88) with the lncRNA Omy200030291 (Additional file 3; Table S14), which positively correlates and interacts with an OMV-specific sRNA *soFE095462* located within the PAI (Additional file 3; Table S13).

In contrast, only 71 upregulated transcripts were identified in resistant fish (Additional file 3; Table S9). Uncharacterized LOC110486996 and anaphase-promoting complex subunit 13 (ANAPC13_1) were the most highly upregulated transcripts. ANAPC13 is involved in cell cycle progression and has been reported to have a potential role in class-I MHC-mediated antigen presentation [36]. ANAPC13_1 exhibited interaction and reciprocal expression with 21 sRNAs (Fig. 6c). Of them, the sRNA *soFE118499* was located in PAI (Additional file 2; Table S6), whereas 20 sRNAs were OMV-specific (Additional file 1; Table S3). Of note, most of the sRNAs were more likely interacting with ANAPC13_1 through lncRNA Omy300040532, which negatively correlates in expression with ANAPC13_1 (Fig. 6c). Interestingly, eight coding transcripts showed contrasting gene expressions (Fig. 6d). These transcripts encode ANAPC13, UTP-glucose-1-phosphate uridylyltransferase, vicilin-like seed storage protein At2g18540, and CCR4-NOT transcription complex subunit 6. Isoforms generated from the same gene locus have previously been reported coding for antagonistic proteins. For instance, the fruit fly *Bcl-x* gene generates two isoforms; one activates apoptosis while the other suppresses it [37].

### Conservation of DE sRNAs in 65 strains with variable degree of virulence

We investigated the conservation of DE sRNAs in 65 strains with variable degrees of virulence. Based on previous studies, 23 *F. psychrophilum* strains were categorized as high virulent strains, and six strains were categorized as low virulent strains (see Methods section) [38–40]. Of 116 DE sRNAs, seven DE sRNAs were conserved in all tested strains, whereas eight DE sRNAs were present in the virulent strains. The list of sRNAs belonging to each group is included in Additional file 3 (Table S15).

Among sRNAs present in virulent strains, a single sRNA (*soFE015758*) was located within a PAI spanning the genomic region 583837-607648. Additionally, the OMV-specific sRNA *soFE030439* was conserved in virulent strains. Whereas sRNAs *soFE031866*, *soFE036061*, and *soFE026739* were enriched in OMVs compared to the whole-cell. sRNA *soFE015758* targets many immune genes, including dedicator of cytokinesis protein 2, leucine-rich repeats and immunoglobulin-like domains protein 1, and tumor necrosis factor-alpha-induced protein 2. Whereas sRNA *soFE030439* strongly interacts with TNF superfamily member 5, myelin-associated glycoprotein, and B-cell receptor CD22.

### *In vitro* treatment of RTgill-W1 cells with OMVs

A cell viability assay was performed to determine if OMVs cause lysis/death in RTgill-W1 cells after different time durations (Fig. 7a). A Presto Blue cell viability assay showed a significant cell lysis/death after OMV exposure of 12 and 24 hours (Fig. 7b). The cytolytic effect of OMVs might be mediated by the enzymes or toxins incorporated in OMVs.

**Fig. 7:**
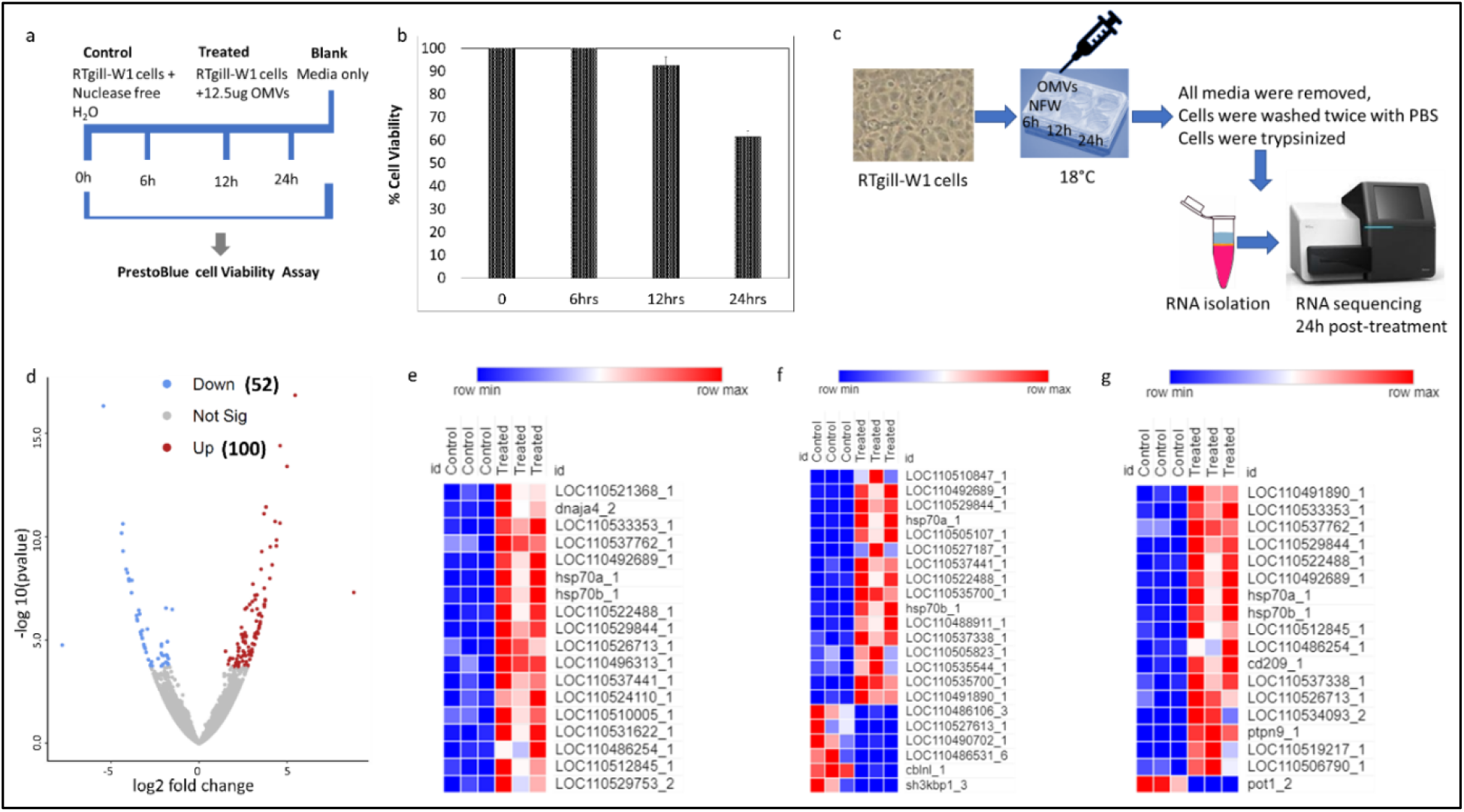
**a)** Overview of PrestoBlue cell viability assay in RTgill-W1 cells after different time durations of OMVs treatment. Approximately 500,000 cells/well were seeded in 96-well plates for 24h. Cells were then treated with OMVs or Nuclease-free water (NFW; control group) for 0 min, 6h, 12h, and 24h at 18°C. Cell culture medium was used as a blank for the PrestoBlue Assay. PrestoBlue reagent was added to wells, followed by incubation at appropriate temperatures and absorbance measurement at 570 nm to determine cell viability. **b)** PrestoBlue cell viability assay in OMVs treated RTgill-W1 cells showed that the cell lysis/death increases with increased time duration of OMVs exposure. Significant cell lysis was observed at 12 and 24h (p < 0.05). **c)** Approximately 1.2 × 10^6^ RTgill-W1 cells/well were seeded in 6-well tissue culture plates and maintained overnight at 18°C. Cells were exposed to either OMVs or NFW (control group) for 6h, 12h, and 24h at 18°C. The media from the “control” and “treated” cell wells were removed, and cells were washed, trypsinized, and then lysed with Trizol reagent for RNA extraction and sequencing on day 1 post-treatment. **d)** A Volcano plot showing the host transcripts DE on day 1 following OMVs treatment in RTgill-W1 cells. The red dots represent the upregulated transcripts in the OMV-treated cells, whereas the blue dots represent the downregulated transcripts at FDR ≤ 0.05. Heat maps showing DE transcripts involved in chaperones and folding catalysts **(e)**, membrane trafficking and endocytic pathway **(f)**, and phagocytosis, and antigen processing and presentation **(g)**.

To further understand the OMV-sRNA interaction with the host genes, we profiled transcriptome expression of RTgill-W1 cells at 24h post-treatment with OMVs (Fig. 7c). RNA sequencing yielded a total of 105,955,286 reads (Average 17,659,214 reads) from six cDNA libraries prepared from control and treated cells (3 libraries/each). A total of 77,160,290 (72.82%) reads were mapped to the rainbow trout genome. 152 trout transcripts were DE in response to OMV treatment (Fig. 7d and Additional file 3; Table S16). Most of them (~66%) were upregulated in treated cells.

Surprisingly, our analysis of the RTgill-W1 cell response to OMV treatment after 24h, considered an early response, did not reveal signs of downregulation of host genes with potential roles in immunity. Only a single gene, complement factor H was downregulated in treated cells (Additional file 3; Table S16). Pathogens exploit complement factor H to inhibit opsonization by C3, and thus resist phagocytosis [41]. Conversely, we observed upregulated expression of several immune-related genes such as complement protein component C7-1, chemokine CXCF1b, TNFAIP3-interacting protein 1, and transcription factor AP-1 (Additional file 3; Table S16). C7 molecule is a constituent of the membrane attack complex (MAC), which forms pores into bacterial target membranes, leading to cell lysis and death [40].

The KEGG analysis provided a general overview of essential genes/pathways regulated following OMV treatment. During the early response to OMVs, genes involved in chaperones and folding catalysts, membrane trafficking and endocytic pathway, phagocytosis, and antigen processing and presentation mainly were upregulated (Fig. 7e-g and Additional file 3; Table S16). Also, 12 transcripts encoding Hsp70, Hsp90, and cochaperones were upregulated in RTgill-W1cells in response to OMV treatment (Fig. 7e).

In addition, some proteins involved in endosomal trafficking were upregulated, including Rab and vacuolar protein sorting proteins. Contrariwise, we noticed downregulation of a few endocytosis-related genes such as PH and SEC7 domain-containing protein 1, protein CLEC16A, and SH3 domain-containing kinase binding protein 1 (sh3kbp1) (Fig. 7f). Fig. 7g shows seven upregulated genes implicated in phagocytosis. The list includes genes essential for pathogen recognition (CD209 molecule), actin remodeling (fibroblast growth factor 7 and transgelin-2), and phagosome acidification (V-type proton ATPase catalytic subunit A; ATP6V1A).

Our results show early inhibition of the suppressor of cytokine signaling 1 at 24h following OMV treatment. *Mycobacterium tuberculosis* induced early expression of a member of the suppressor of cytokine signaling (SOCS) family of proteins to control phagosomal acidification by selectively targeting the ATP6V1A for degradation [42]. We also noticed downregulation of a transcript encoding protection of telomeres 1 (POT1), which negatively regulates phagocytosis. Additionally, ten upregulated genes involved in antigen processing and presentation were identified. These genes encode beta-2-microglobulin (B2M) and nine heat shock proteins (Fig. 7g).

Furthermore, genes with anti-apoptotic roles, such as AP-1 proteins, DNA-damage-inducible transcript 4 (DDIT4), heat shock proteins (Hsp70 and Hsp90), and cochaperones (DnaJ/Hsp40 and tetratricopeptide repeat (TPR)), were upregulated (Additional file 3; Table S16). However, a few genes whose products likely have apoptotic properties, such as MAP/microtubule affinity-regulating kinase 4 and Growth arrest and DNA-damage-inducible protein GADD45 beta, exhibited contrasting expression (Additional file 3; Table S16).

Notably, 69 bacterial sRNAs showed significant expression (>100 reads) in the treated cells. Of them, 13 sRNAs belong to the cell-enriched or OMV-enriched/-specific categories (Table 3). The top four sRNAs targeted most of the downregulated host transcripts following OMV treatment. For instance, the OMV-enriched sRNA *soFE013584* exhibited a very high interaction (ndG = −0.54) with the transcript encoding POT1. Also, the OMV-enriched sRNA *soFE002123* interacted with the POT1. POT1 protein is vital for the proper maintenance of genome integrity. Disruption of POT1 function results in chromosome instability and loss of cellular viability [43].

**Table 3:** OMV-enriched/-specific versus whole cell-enriched bacterial sRNA detected 24h following treatment of the RTgill-W1 cells with OMVs. The negative log2 ratio indicates that sRNA is less abundant in OMV than the whole-cell.

| ID | Start | End | Read counts | Category (log<sub>2</sub> ratio) | PAI |
| --- | --- | --- | --- | --- | --- |
| soFE125269 | 1637553 | 1637893 | 3145 | cell-enriched (-1.72) | No |
| soFE004028 | 2056951 | 2057269 | 2062 | OMV-specific | No |
| soFE013584 | 2196235 | 2196334 | 1912 | OMV-enriched (3.1) | No |
| soFE002123 | 2061751 | 2061850 | 1020 | OMV-enriched (4.23) | No |
| soFE104747 | 1635059 | 1635287 | 918 | OMV-specific | No |
| soFE048173 | 2195010 | 2195109 | 657 | OMV-enriched (4.21) | No |
| soFE011466 | 597087 | 597258 | 272 | cell-enriched (-1.43) | Yes |
| soFE091624 | 595998 | 596220 | 264 | OMV-enriched (1.61) | Yes |
| soFE004713 | 1634302 | 1634689 | 231 | OMV-specific | No |
| soFE100359 | 593374 | 593473 | 216 | cell-enriched (-1.31) | Yes |
| soFE073471 | 2540111 | 2540210 | 210 | OMV-enriched (1.63) | No |
| soFE011096 | 2046508 | 2046607 | 181 | cell-enriched (-1.2) | No |
| soFE100358 | 593129 | 593228 | 133 | OMV-specific | Yes |

## Discussion

Several studies have focused on the role of eukaryotic membrane vesicles carrying nucleic acid content (DNA, RNA, miRNA) in disease states [44]. However, less is known about the role of nucleic acids present within prokaryotic membrane vesicles in affecting the host during the host-pathogen interactions. Previous studies have demonstrated the presence of DNA, peptidoglycan, and lipopolysaccharides (LPS) in OMVs purified from bacterial cultures [45, 46]. These cargos can be delivered to host cells by binding OMVs to surface receptors on host cells or through endocytosis [47, 48]. Because OMVs provoke the transfer of various inner constituents from bacteria to the host, it has driven our attention to further investigate the effect of OMVs on host-pathogen interactions. Thus far, only a few studies [16, 49] have associated the presence of sRNAs within bacterial OMVs with a role in host health. Dual RNA-seq identified sRNAs produced from *Salmonella enterica*, such as *PinT*, demonstrated a role in regulating the expression of host genes and mediating the activity of virulence genes necessary for intracellular survival of the pathogen [50]. Koeppen *et al*. reported that *P. aeruginosa* sRNAs target kinases in the LPS-stimulated MAPK signaling pathway [16]. sRNAs in the plant pathogen *Xanthomonas compestris pv vesicatoria* were also involved in pathogenicity, and deletion of the sRNA led to a reduction in virulence of bacteria [51].

In this study, RNA-Seq analysis revealed the presence of sRNAs in OMVs and found variation in sRNA abundance between OMV and whole-cell transcriptome. Our results indicated that OMV-specific/-enriched sRNAs target immune-relevant genes. These sRNAs included *soFE129980* (TPM 830), which targets the host NLR family CARD domain-containing protein 3; *soFE081095* which targets butyrophilin subfamily 1 member A1; and sRNA *soFE023505*, which targets interferon-induced protein 44 and major histocompatibility complex class I-related gene protein. Similar to our work, a recent study used RNA-Seq to characterize OMVs-packaged sRNAs in *P. aeruginosa* and predicted their human immune transcript target. The study reported a specific OMV-sRNA delivered to host cells, resulting in OMV-induced attenuation of IL-8 secretion and neutrophil infiltration in mouse lungs [16].

Although many attempts have been made to understand the mechanism of host-pathogen interactions between rainbow trout and *F. psychrophilum* [6–8], very little is known regarding the pathogenesis of this bacterium. Differential expression of trout genes between the ARS-Fp-R and ARS-Fp-S genetic lines has been previously reported [6]. However, the potential influence of the bacterial sRNAs in regulating the trout genes and their potential effect on the survivability of the fish was not previously reported in rainbow trout. Our study showed that most DE sRNAs were upregulated in fish susceptible to BCWD. OMV-enriched sRNAs exhibited strong interactions and differential reciprocal expression with trout genes in BCWD-resistant and -susceptible genetic lines. sRNAs interacting with the trout genes share complementary sequences, which ultimately might cause post-transcriptional repression of the trout genes. sRNAs targeting immune genes might help the bacteria to highjack the immune response and modulate the survival of bacteria in the host [52]. The effect of sRNAs was tested in *Staphylococcus aureus* by silencing the sRNA, which modulated the virulence and disease resistance of *S. aureus* [53]. Similarly, another study in *Staphylococcus* reported the role of sRNA in bacterial virulence and regulating the expression level of an immune evasion molecule [54]. Further investigation beyond the scope of this study is required to determine whether deletion of specific bacterial sRNAs can affect the *F. psychrophilum* invasion of the host. Identifying new microbial targets may facilitate controlling bacterial infections and containing the BCWD.

The study revealed a significant cell lysis/death at 12 and 24 hours following *in vitro* exposure of RTgill-W1 cells to 12.5 μg/mL OMVs. Consistently, *Burkholderia cepacia* OMVs induced cytotoxicity in host cells treated with ≥10 μg/mL of OMVs for 24h [55]. In addition, OMVs induced cytotoxicity of the salmonid CHSE-214 cells [13]. Most of the DE transcripts involved in cell death regulation are encoding products with anti-apoptotic roles, suggesting that cell death occurred by necrosis due to exposure to OMV toxins. One characteristic of the OMV treatment is the upregulation of transcripts for HSP70, HSP90, and cochaperones. Cochaperones, such as DnaJ/Hsp40 and tetratricopeptide repeat (TPR), determine the multifunctional properties of Hsp70 and Hsp90, including their anti-apoptotic functions [56]. The virulence factor Hsp60 was identified in the proteome of OMVs of the fish pathogen *Piscirickettsia salmonis[13]*. In addition to their role in regulating cell death/apoptosis, heat shock proteins stimulate elements of innate immunity and traffic antigens into antigen-presenting cells, which facilitates induction of specific acquired immune responses (reviewed in [57]).

Moreover, OMV treatment modulated host gene expression, favoring elements from the phagocytic, endocytic, and antigen presentation pathways. Phagocytosis is a conserved defense strategy in which host cells engulf and destroy self/nonself antigens. Although the epithelial RTgill-W1 cells are not professional phagocytes, epithelial cells were previously reported to have low phagocytic activity and a significant contribution to pathogen clearance [58]. Following OMV treatment, RTgill-W1 cells upregulated the phagocytic receptor CD209 (DC-SIGN) responsible for pathogen recognition [59]. Upregulation of genes involved in the regulation of actin cytoskeleton, such as fibroblast growth factor and Transgelin-2, suggests that actin remodeling is essential at the site of phagocytosis. It has been previously reported that Transgelin-2 [60] and fibroblast growth factors [61, 62] enhance phagocytosis. Several other genes involved in phagocytosis, including ATP6V1A, were also upregulated. V-ATPase molecules accumulate on the phagosomal membrane to acidify the phagosome interior [63, 64]. Some pathogenic bacteria (*M. tuberculosis*) have developed a strategy to survive inside host phagocytes by facilitating ATP6V1A degradation [42]. In addition to phagocytosis, genes implicated in the endocytic pathway, such as early endosome antigen 1 (EAA1), vacuolar sorting proteins, and Rab proteins, were also regulated in response to OMV treatment. EAA1 is recruited to the phagosome, promoting early endosomal fusion with the new phagosome [65].

Moreover, OMV-treated cells upregulated B2M, a scaffolding protein that retains the native structure of MHC class I molecules on the surface of nucleated cells to present antigens to cytotoxic CD8+ T cells. In addition to the role of B2M in adaptive immunity, it possesses antimicrobial activities (i.e., innate response). B2M sheds sB2M-9 fragments, which function as antibacterial chemokines, and perhaps as potential antimicrobial peptides (AMPs) following modification by thioredoxin [66]. Upregulating the expression of phagocytosis-, endocytosis-, and antigen presentation-related genes reflect host attempts to increase uptake of OMVs and present processed antigens to stimulate the immune response.

OMVs possess some inherent characteristics that qualify them as vaccine candidates (reviewed in [67]). For instance, OMVs are storage containers that can remain intact under different treatments and temperatures. Also, OMVs have non-living activity, adjuvant effects, non-replicative properties, and hold substantial immunogenic components belonging to the parent bacteria, which may induce immune responses against bacterial infection. OMVs derived from *Salmonella* Enteritidis [68] and *Bordetella bronchiseptica* [69] provided substantial protection against the parent bacterial infection. However, further research is still needed to study the role of *F. psychrophilum*-derived OMVs in boosting the host immune response, and thus their potential application as a vaccine to contain/control the BCWD.

## Conclusions

The current study provided the first characterization of *F. psychrophilum* sRNAs in OMVs and demonstrated their potential effects on host gene expression *in vivo and in vitro*. One key aspect of this study is the simultaneous capture of the RNA expression profile of the *F*. *psychrophilum* and its piscine host (rainbow trout) following infection in selectively bred resistant and -susceptible genetic lines. The whole-body dual RNA-seq approach yielded a comprehensive transcriptomic dataset that enabled us to identify microbial factors contributing to the disease progression at a later stage of the infection. In particular, we identified bacterial sRNAs packaged/enriched within OMVs and delivered to the host. These sRNAs revealed strong interactions and reciprocal expression profiles with the host immune genes. Such a relationship suggests a role for the OMV-enriched sRNAs in shaping the host-pathogen interactions and assisting the bacteria to highjack the host immune response. Future studies considering the early and late stages of infection will provide insights into the dynamics of gene expression changes during infection.

Additionally, the study revealed significant cell lysis/death at 12 and 24 hours following *in vitro* exposure of RTgill-W1 cells to OMVs. The OMV treatment also modulated host gene expression, favoring elements from the phagocytic, endocytic, and antigen presentation pathways. Further studies are needed to investigate whether OMVs can induce immune memory in rainbow trout and thus assess their potential use as a vaccine.

## Materials and Methods

### Ethic Statement

Fish were maintained at the NCCCWA and animal procedures were performed under the guidelines of NCCCWA Institutional Animal Care and Use Committee Protocols #053 and #076.

### Bacterial culture

*F. psychrophilum* (CSF 259-93) used in this study was obtained from USDA/NCCCWA (Dr. Gregory Wiens). Frozen stock cultures of *F. psychrophilum* were cultured on Tryptone Yeast Extracts (TYEs) agar, and the plate was incubated at 15°C for one week. *F. psychrophilum* colonies isolated from the TYEs agar plate were transferred to TYEs broth, and absorbance (525nm) was measured after 24h of incubation. Absorbance was measured every day for 2 weeks to determine the log phase of the cultures. TYEs broth culture without *F. psychrophilum* was used as a negative control.

### Isolation of OMVs

OMVs were isolated from *F. psychrophilum* broth culture on day 8 of bacterial growth. A loopful of culture was sub-cultured onto a plate on day 7 of bacterial growth to ensure the broth was free of contamination. For OMV isolation, broth culture from a flask was distributed into several 50 ml tubes. Each tube was centrifuged at 2800 X g for 1h at 4°C to pellet the bacterial cells. The supernatant was collected and filtered through a 250 ml sterile 0.22μm PES membrane filter (EMD Millipore Corporation, Billerica, MA, USA) to remove any remaining bacterial cells. The filtrate was then subjected to ultracentrifugation for 3h at 40,000 rpm and 4°C to pellet the OMVs. The OMV pellet was washed with phosphate-buffered saline (PBS) and again subjected to ultracentrifugation for 2h at 40,000 rpm and 4°C to re-pellet the OMVs. The OMV pellet was resuspended in nuclease-free water and stored at −20°C. The protein concentration of the OMVs was quantified using a BCA Protein assay kit (Thermo Fisher Scientific). To ensure that the suspension containing OMVs was free of bacteria, 30 μl of the suspension was cultured on a TYEs agar plate and incubated for 10 days.

### Transmission Electron Microscopy (TEM)

Transmission electron microscopy was performed on *F. psychrophilum* cell and OMV samples. For *F. psychrophilum* cells, a single colony from a TYEs-agar plate was suspended in nuclease-free water. Water and TYEs broth were used as negative controls. All samples were subjected to negative staining using uranyl acetate, and samples were deposited on a TEM Carbon coated grids (80 mesh square grid, EMS; Ted Pella, Inc., Redding, CA, USA) and incubated for 2 min. Samples were blotted dry, and grids were washed with sterile deionized water (diH2O) three times (30 seconds each) to remove the salt buffer. Excess water from grids was removed with blotting paper before the samples were stained. Samples were stained with 5 μl of 1% uranyl acetate added onto the grid and incubated for about 1 min. The stain was then washed with sterile diH2O, dried, and the grids were observed under a Hitachi H-7650-II (Schaumburg, IL, USA) transmission electron microscope.

### Bacterial RNA extraction, cDNA Library Preparation, and Sequencing

RNA was extracted from *F. psychrophilum* colonies isolated from a TYEs agar plate and from OMVs isolated from *F. psychrophilum* broth culture using TriZol reagent (Invitrogen, Carslsbad, CA, USA). The RNA concentration was measured using a Nanodrop™ ND-1000 spectrophotometer (Thermo Fisher Scientific, Waltham, MA, USA). To confirm the existence of RNA in OMVs, 4 μl of the RNA samples were treated with 2 μg/ul of RNase Cocktail Enzyme mix (Thermo Fisher Scientific Baltics, VA) and incubated in a water bath at 37°C for 30 min. RNase-treated and -untreated OMV RNA samples were then run on an agarose gel. RNA samples isolated from *F. psychrophilum* and OMVs were stored at −80°C until further processing.

For library preparation and RNA sequencing, RNA samples isolated from *F. psychrophilum* cells and OMVs were sent to BGI Genomics (Cambridge, MA, USA). Library preparation was performed using a Trio RNA-Seq kit (NuGEN, San Carlos, CA, USA) according to manufacturer recommendations. Briefly, rRNA was depleted and RNAs were fragmented using a fragmentation buffer. Fragmented pieces were then purified using QiaQuick PCR extraction kit (QIAGEN, Germantown, MD, USA), and the solution was resuspended in EB buffer, and cDNA was subjected to end repair and poly (A) tail addition. The fragments were then connected with the adaptors. The library was then subjected to purification using a MiniElute PCR Purification kit before PCR amplification. PCR was used to amplify the libraries, and then the yield was quantified. Sequencing was performed on an Illumina MiSeq platform (Illumina, Inc., San Diego, CA, USA). Raw RNA-Req reads were submitted to the NCBI Short Read Archive under accession number BioProject ID PRJNA259860 (accession number SUB10837562).

### Prediction of sRNAs and their target genes

Raw sequence reads were trimmed by removing adaptor sequences and 5 bp from each end, followed by filtering out low-quality reads. High-quality reads from *F. psychrophilum* (60,352,578 reads) and OMVs (55,722,742 reads) were used for downstream analyses. Reads from the *F. psychrophilum* transcriptome and OMVs were mapped to the *F. psychrophilum* (CSF-259-93) reference genome using TopHat2 [70]. The FlaiMapper tool [28] was used to identify potential sRNAs. sRNAs were predicted and filtered according to their length. sRNAs greater than 500nt long were filtered out. DE transcripts and trout immune-related genes targeted by the DE sRNAs were predicted using a locally installed LncTar software [71]. A normalized deltaG (ndG) of −0.10 was used as a cutoff value to determine the potential interaction among paired RNAs.

### Fluorescent in situ hybridization (FISH) and Confocal Microscopy

FISH staining was used to confirm the presence of sRNAs within *F. psychrophilum*. Briefly, *F. psychrophilum* cultures from the log phase were harvested and centrifuged at 2800 rpm for 5 min at 4°C. The supernatant was discarded and the pellet containing the cells was washed with PBS and then fixed with 4% paraformaldehyde in PBS for 4h at 4°C. After fixation, 10 μl of the cells were smeared onto a microscopic glass slide, and the smear was air-dried. Cells were then dehydrated in an alcohol series (50, 80, and 100% ethanol) for 3 min in each concentration at room temperature. Cells were then hybridized with a Cy3 fluorophore-conjugated sRNA probe (*soFE128978*) for 2h at 46°C in the dark. A non-sense eubacterial probe and a sense eubacterial probe with FAM fluorophore were used as negative and positive controls. To prevent drying, slides were incubated in a hybridization chamber. After hybridization, cells were washed and then incubated with washing buffer (5M NaCl, 1M Tris/HCl, 0.5 M EDTA, 10% SDS and distilled H2O) at 48°C for 25 min. Then cells were incubated for 3 min in PBS containing DAPI (300 nM). DAPI-stained slides were then washed with water and air-dried. Glass coverslips were mounted to each slide after adding a drop of VectaShield (Vector Laboratories, Burlingame, CA, USA); the edges of the coverslip were sealed with clear nail polish. Once dried, the slides were observed using a Zeiss Axio Observer microscope with LSM700 confocal module. Imaging was performed with ZEN 2012 software (Black Edition, Carl Zeiss Microscopy, Thornwood, NY, USA). Smart Setup function of the ZEN 2012 imaging software was used to assign the optimal filter and beam splitter settings for each laser while laser power and gain were set manually.

### BCWD-resistant and -susceptible fish population, and bacterial challenge

The current study used samples from two rainbow trout genetic lines of divergent resistance to BCWD. USDA-NCCCWA produced these genetic lines via a family-based selection method as previously described [72]. Briefly, single-sire × single-dam matings were made between 3-year-old dams and 1-year old sires (neo-males) within the genetic lines. The resistant line eggs were produced from dams that had undergone three generations of BCWD selection, whereas the sires had undergone four generations of selection to increase the disease resistance phenotype. On the other hand, the susceptible line eggs were produced from parents that had undergone one generation of selection to increase susceptibility to infection. The resistant and susceptible genetic lines had significant differences in susceptibility to F. psychrophilum [6, 7].

For this study, fish from resistant and susceptible genetic lines (49 days post-hatch) were challenged with F. psychrophilum as we previously described [6]. Briefly, fifty fish were randomly assigned to each tank with 2.4 L min−1 of 12.5 ± 0.1 °C flow-through spring water supply. For each genetic line, two fish tanks were intraperitoneally injected with 4.2 × 106 CFU fish−1 F. psychrophilum suspended in 10 μl PBS, and survival was monitored daily for 21 days. Five individuals were sampled from each tank on day 5 post-injection. All fish were certified to be infection-free before being injected with F. psychrophilum.

### RNA sequencing and gene expression analysis of BCWD-resistant and -susceptible fish

As explained above, tissue samples used in this study were obtained from USDA/NCCCWA (Dr. Gregory D Wiens). RNA was extracted from the whole fish using TriZol, followed by quantity and quality assessments as described above. RNA samples were then treated with DNAase I (Fisher BioReagents, Hudson, NH, USA) to remove DNA from samples. For sequencing, equal amounts of RNA samples were pooled from 2 fish, and 4 pooled samples were sequenced from each of the resistant and susceptible genetic lines (i.e., a total of 8 libraries). Sequencing was done at RealSeq Biosciences, Inc. (Santa Cruz, CA, USA). The Zymo Ribofree library prep kit targeted all RNAs during the library preparation, including trout and bacterial RNAs.

Raw sequence reads from each genetic line were trimmed to obtain high-quality reads. Read mapping and gene expression analyses were performed using a CLC genomics workbench. Reads were separately mapped to the rainbow trout genome [USDA OmykA_1.1 assembly (GCF_013265735.2)] and F. psychrophilum reference genomes to identify DE transcripts. Mapping criteria were as follows: mismatch cost = 2, insertion/deletion cost = 3, minimum length fraction = 0.9, and similarity fraction = 0.9. The expression value of each transcript was calculated in terms of TPM and DE transcripts between resistant and susceptible genetic lines were identified using EDGE test (FDR value < 0.05, log2 fold change cutoff ±1). Raw RNA-Req reads were submitted to the NCBI Short Read Archive under accession number BioProject ID PRJNA259860 (accession number SUB10837562).

### Maintenance of RTgill-W1 cell line

Frozen RTgill-W1(CRL-2523) cells were obtained from ATCC (American Type Culture Collection, Manassas, VA, USA). Cells were handled according to instructions provided by ATCC. Briefly, frozen cells were thawed at 20°C in a water bath, and the full content from the vial was transferred to L-15 media supplemented with 10% complement inactivated fetal bovine serum (FBS) and 100 units/ml of penicillin and 100 μg/ml of streptomycin (complete medium). Cells were centrifuged at 125 X g for 8 min, the supernatant was removed, and the cells were resuspended in a complete medium. The cells were maintained in 25 cm2 flasks at 18°C. Media were renewed every 4 days, and cells were passaged every 8 days. Since cells are adherent, cells were dislodged by trypsinization while passaging.

### Exposure of RTgill-W1 cells to OMVs

Approximately 1.2 × 10^6^ RTgill-W1 cells/well in L-15 media supplemented with 2% FBS [73] were seeded in 6-well tissue culture plates. Before treatment, cells were maintained overnight at 18°C. A 100 μl volume of OMVs (total protein concentration = 0.8 mg) was added to cells for 6h, 12h, and 24h. For a control, nuclease-free water was added to cells. Each condition was run in triplicate. Plates were incubated at 18°C, and sampling was done after each incubation period. All the media from the “control” and “treated cells” were removed, and cells were then washed twice with PBS. The cells were trypsinized and then lysed with Trizol reagent (Invitrogen, Carslsbad, CA, USA) for RNA extraction, followed by sequencing and gene expression analyses. To determine the cell viability, we performed a PrestoBlue Assay in which ~500,000 cells/well were seeded in a 96 well plate for 24h. Cells were then treated with OMVs for different time intervals of 0 min, 6h, 12h, and 24h. Each condition was run in triplicate. Nuclease-free water was added to cells as a control, and only cell culture medium was used as a blank for the PrestoBlue Assay. After treatment, PrestoBlue reagent was added to wells, and then the plate was incubated at 37°C for 10 min followed by incubation at 18°C for 1h. The absorbance of treated cells and controls was then measured at 570 nm to determine cell viability.

### sRNAs conservation and identification of sRNAs within pathogenicity Islands

To determine the conservation of sRNAs, DE sRNAs reported in this study were blasted against the genome sequences of 65 different *F. psychrophilum* strains downloaded from the National Center for Biotechnology Information (NCBI) GenBank database (https://www.ncbi.nlm.nih.gov/genome/browse/#!/prokaryotes/1589/). *F. psychrophilum* strains used in this study were 950106-1/1, DSM 3660, 010418-2/1, 4, 10, 11754, 17830, 160401-1/5, 990512-1/2A, CH8, CH1895, CHN6, CN, CR, F164, FI055, FI070, FPG3, FPG48, FPRT1, FPG1, FPS-F15, 160401-1/5N, P30-2B/09, F164, FPS-F16, FPS-S11A, 990512-1/2A, P15-8B/11, 160401-1/5M, FPS-F27, FPS-R7, FPS-S9, JIP02-86, FPS-S6, OSU THCO2-90, FPS-P3, FPS-P1, V46, JIP 08/99, JIP 16/00, FPS-S10, 010418-2/1, K9/00, 950106-1/1, FPG3, FPGIW08, IT02, IT09, IWL08, K9/00, KKOK-1706, KU060626-59, LM01-Fp, MH1, NNS1-1804, NNS4-1804, NO014, OSU-THCO2-90, P7-7B/10, P158B-11, P30-2B, TN, TR). The high virulent Fp strains included FPG1, FPS-F15, 160401-1/5N, P30-2B/09, F164, FPS-F16, FPS-S11A, 990512-1/2A, P15-8B/11, 160401-1/5M, FPS-F27, FPS-R7, FPS-S9, JIP02-86, FPS-S6, OSU THCO2-90, FPS-P3, FPS-R9, FPS-P1, V46, JIP 08/99, JIP 16/00) [40], whereas less virulent Fp strains included FPS-S10, 010418-2/1, K9/00, 950106-1/1, FPG3, and FPGIW08 [40, 74]. A threshold of >95% identity and 100% query coverage were used as the cutoff values for homology.

The webserver IslandViewer 4 (Integrated Interface for Computational Identification and Visualizations of Genomic Island; http://www.pathogenomics.sfu.ca/islandviewer/) [75] was used to identify and visualize pathogenicity islands present within the *F. psychrophilum* genome sequence. Bedtools [76] was used to determine the location of sRNAs within the islands.

